# Sex-specific effects of developmental density on life history and age-dependent flight traits

**DOI:** 10.1101/2020.08.24.265736

**Authors:** Krish Sanghvi, Felix Zajitschek, Maider Iglesias-Carrasco, Megan L. Head

## Abstract

The environment organisms experience during development can have effects which carry over into their adult lives. These developmental environments may not only affect adult traits at a given point in time, but also how these traits change with age. Generally, stressful developmental environments can lead to sub-optimal adult fitness traits and a faster deterioration of these traits with age. But whether these environments affect how performance traits change with age or whether they affect males and females differently lacks evidence. Here, we test the sex-specific effects of crowding during development on life history traits such as lifespan, fecundity, age-dependent survival, and reproductive senescence, as well as flight performance traits, in the seed beetle-*Callosobruchus maculatus*. We show that although developmental crowding has no effect on flight performance of either sex, on male adult lifespan, or on female age-dependant survival, it is detrimental to female lifespan and fecundity, as well as female age-dependent reproduction. Additionally, we show that flight traits can change with age and differ between males and females, which may be reflective of sex-specific life-histories and behaviour. Our study has implications for understanding how developmental environments affect life history and behavioural strategies of both males and females.

## Introduction

An individual’s developmental environment can have long lasting effects on its phenotype, which carry over into its adult life (English et al, 2016; Lee et al, 2013). For instance, empirical studies often show that individuals who experience stressful environments (e.g. extreme temperatures or poor nutrition) early in life, perform worse as adults in comparison to those who do not (Jonsson & Jonsson, 2014; Lummaa & Clutton-Brock, 2002; Monaghan, 2008). Examples include organisms from across the animal kingdom, in which individuals who developed in stressful environments had shorter lifespans as adults (van de Pol et al, 2006), were less fecund (Cam & Hines, 2003), had poorer performance (Fleming et al, 1997), and faster deterioration of traits (Nussey et al, 2007) than those from optimum developmental environments.

The population density that individuals experience during development is one of the environmental parameters which has received considerable attention in the context of how developmental environments affect adult traits. There is a general consensus that high density during development has negative effects on individual physiological and life history traits (reviewed in Peters & Barbosa, 1977). For instance, animals which develop under crowded conditions have a smaller body size (Flockhart et al, 2012; Yang et al, 2015), lower reproductive output (Muller et al, 2016; Prout & McChesney, 1985; Scott, 1994), and shorter lifespans (Descamps et al, 2008; Fantinou et al, 2008). However, the effects of high developmental density have been shown to be beneficial for other traits which may also be important in determining fitness. For instance, experiencing greater developmental density increases dispersal ability (Applebaum & Heifetz, 1999; Braendle et al, 2006), competitiveness (Stockley & Seal, 2001), disease resistance (Wilson et al, 2002), and development rate (Bauerfeind & Fischer, 2005). Despite the considerable interest in how developmental environments affect adult traits, there is little research that tests whether the developmental environment influences how adult traits change with age.

Measuring the effects of developmental environments may be particularly important for behavioural, physiological, and life history traits such as reproduction, locomotion, and performance, which are labile rather than static. This is because early life environments may not only affect these traits at a given point in time, but may also affect how these traits change throughout an individual’s lifetime (Balbontin & Moller, 2015). For instance, senescence, which is the intrinsic deterioration of fitness components with increasing age, has been shown to accelerate when early life environments are stressful (Cooper & Kruuk, 2018). Studies show that age-dependent survival is generally lower for those individuals reared at high densities than for those reared in low densities (Graves & Mueller, 1993; Joshi & Mueller, 1997). Effects of developmental densities have also been shown on age-dependent reproduction. For instance, in *Drosophila*, a study shows that individuals from high developmental densities have a later peak and slower decline in reproduction with age (Lints & Lints, 1969).

Performance traits which are ecologically important but are not direct measures of fitness may also be expected to deteriorate with age. For example, studies by Collatz & Wilps (1986) and Ahman & Karlsson (2009) have shown that insect flight deteriorates with age. However, more evidence is needed to test the generality of these results across species and across environments. Studies which test the effects of developmental environments on age-dependent flight traits are especially important because if dispersal ability declines with age at different rates in different environments, the developmental environment will influence the decision to disperse and the optimum timing of dispersal. Testing this will help us understand the different strategies individuals use to avoid crowding and find mates, and help us predict how individuals invade novel environments (Ochocki & Miller, 2017). Additionally, if deterioration of flight traits with age differs between males and females, it may have important implications for sex-biased dispersal and avoidance of inbreeding (Pusey, 1987).

Here we test whether developmental density affects not only adult life history traits (i.e survival and fecundity) but also flight performance in the seed beetle *Callosobruchus maculatus* (J. Fabr. 1775), and how these traits change with age. Previous studies have shown that seed beetles exhibit strong responses to density, with high developmental densities causing individuals to have shorter developmental times, longer lifespans, and lower fecundity (Vamosi, 2005). Some studies have also suggested that seed beetles reared in high developmental densities may produce a dispersal morph which has greater flight ability in order to disperse away from crowded conditions (Appleby & Credland, 2001; Messina & Renwick, 1985; Utida, 1972), although previous tests for measuring flying ability are not convincing due to the lack of repeatable methodology (see Appendix S1). Furthermore, how adult life history and performance traits change with age under different developmental densities, has to our knowledge not been tested in seed beetles before.

## Methods

### Material

Our stock population of *C. maculatus* was originally sourced from stock kept at the University of Western Australia since 2005 (Dougherty et al, 2017). Prior to the current experiment, this stock was raised in our lab for 9 generations on cowpeas (*Vigna unguiculata*) under crowded conditions (approximately between 10-15 eggs laid on each bean and 200 beans per container), at 24°C to 28°C.

### Experimental design

To investigate how developmental density affects adult life history and flight performance traits, and the way these traits change with age, we set up two developmental densities: low and high.

For the low developmental density treatments, 45 females were randomly chosen from our stock population within 3 days of their eclosion, and were individually placed in a petri dish (10 cm diameter), along with two males and 15 cowpeas. Females and males were left overnight in these dishes to mate and lay eggs on the cowpeas. The following morning the cowpeas were checked for eggs, and 200 cowpeas out of the total 675 beans, were randomly chosen from across the 45 dishes. If the bean had more than one egg laid on it, the extra eggs were scraped off prior to being used to ensure that all beans used had only one egg on each of them Cowpeas were then transferred into five cylindrical containers (2 cm x 4.5 cm) which had holes in the lid for ventilation, with each container holding 40 beans.

For the high developmental density treatment, 150 females and 300 males were randomly chosen from our stock population within 3 days of their eclosion. A total of five female and ten males were placed in each petri dish (10 cm diameter) with 15 cowpeas per dish, and a total of 30 such petri-dish replicates were set up. Females and males were left overnight in these dishes to mate and lay eggs on the cowpeas. The following morning, the cowpeas were checked for eggs and a total of 200 cowpeas, each with at least five eggs laid on them, were randomly sampled from across the 30 petri dishes (200 beans sampled out of 450). Forty cowpeas were transferred into each of five cylindrical containers (2 cm x 4.5 cm), which had holes in the lid for ventilation. In the high density treatment, the number of eggs per container ranged between 216 and 280. For both treatments, all eggs in each container were laid on the same day and were incubated between 27 °C to 28 °C, and between 20% and 40% relative humidity.

### Development time and survival

We checked the containers daily for emerging beetles. As adults emerged, we recorded their development time (time elapsed between date of laying and date of eclosion) and transferred them to individual Eppendorf tubes with a small hole in the lid for ventilation, with each tube being given an individual label. Because *C. maculatus* adults do not need to feed or drink after eclosion, we did not provide them with food or water. The sex of emerged beetles was identified following Beck & Blumer (2011). Once in individual tubes, we checked adult beetles daily to determine whether they were dead or alive, so that we could calculate their adult lifespan (days elapsed between emergence and death). A total of 501 females and 535 males emerged from the high-density treatment, while a total of 85 females and 100 males emerged from the low-density treatment, all of which had their lifespans measured.

### Female fecundity

To determine the number of eggs laid each day throughout a female’s lifetime and her total fecundity, we selected a random subset of females from each density treatment (High density: N= 69, Low density: N= 49) on the same day that they eclosed, and transferred them into individual petri dishes with 2 random males from the stock population and 15 cowpeas, which were replaced daily with a new set of 15 cowpeas. After the cowpeas had been laid on, they were stored in small plastic bags in a −20 °C freezer, so that the eggs could be counted later. We sampled fewer females from the low density treatment because only a total of 85 females emerged from this treatment, and we wanted to balance the number of low-density females across the virgin-flight assay (see below) and fecundity assay.

### Measuring flight

We measured flying ability and propensity to fly for 86 virgin females and 162 virgin males from the high-density treatment, and 32 virgin females and 98 virgin males from the low-density treatments, every second day, from the day they eclosed until 18 days after emergence. In addition, we measured the flight traits of all the females which were used in the female fecundity assays (mated females (N): High density=69, Low density=49), in the same way as the virgin-flight assay to test the effect of mating on flight. To elicit a flight response, we probed beetles by using the back of a pair of soft forceps to tap on the beetle’s body and then observed the beetles for 15 seconds. We repeated this process and recorded the beetle’s flight ability and propensity to fly across the whole 30 seconds.

Flying ability for each individual was scored using an ordinal point system based on the action they performed during the trial. Scores ranged from 0 to 3. Beetles which only crawled and showed no wing movement, received a score of “0”. Beetles that opened their wings but did not leave the ground received a score of “1”. Beetles that opened their wings and left the ground but did not fully fly received a score of “2”. Finally, beetles that opened their wings and flew fully in a vertical direction received a score of “3”. We compared our scoring system to that used by Utida (1972) and Appleby & Credland (2001), which drops individuals from a height and gives them a binomial score based on whether they fly or not. We found the method of probing individuals to elicit a flight response to have fewer false negatives and be more repeatable than expected by chance (see appendix S1), hence used this method for our experiment. A similar method which probes individuals to elicit flight has been previously used effectively by Esterhuisen et al (2014) and Frazier et al (2008) in Diptera.

Flying propensity, on the other hand, was the number of times a beetle performed the action of its flying ability. This also included the number of times a beetle flew fully after having landed on a surface. Beetles which did not perform any action (with an ability score of zero), were excluded from this analysis. All flight measures were scored blind to the density treatment the beetle came from.

### Data analysis

#### Age-independent effects

To test whether developmental density affected proportion of beetles that emerged from eggs or affected the sex ratio of eclosed beetles, we ran two separate logistical regression model using the function *glm* in package *stats* in R v3.5.2 (R Development Core Team, 2011). Both models used a binomial error distribution (family=binomial, “logit” link function), and included developmental density as a fixed effect. To model the effects of density on emergence of beetles, we used the binomial variable, emergence (yes/no), as the dependent variable. To model the effect of density on sex ratio, we used the binomial variable, sex of eclosed beetles (male/female), as our dependent variable. We checked both the models for over-dispersion using the package *DHARMa* in R v3.5.2 (Hartig, 2020), and they were not over-dispersed (P>0.9 for both). Neither model used container as a random effect because container was our level of replication. To analyse the effects of developmental density on male and female static traits (i.e. development time, lifespan) we used linear mixed models (LMM). In both models, developmental density, beetle sex, as well as their interaction were included as fixed effects while the container of beans that a beetle came from was included as a random effect. To look at the effect of density and sex on lifespan, we only modelled data from virgin beetles (i.e. we excluded females used in the fecundity assays) in order to avoid the known effects of female mating on adult traits (Jigisha et al, 2020; Ronn et al, 2006). However, to explicitly test for effects of female mating status and its interaction with density on female adult lifespan, we ran a second LMM in which female adult lifespan was the response variable, and female mating status, developmental density, and their interaction were specified as fixed effects. In this model, container was again specified as a random effect.

For all analyses, we ran a “full” model with the interactions wherever present, as well as a “main-effects” model which excluded the interaction and allowed interpretation of the independent effects. For all models, residuals were checked visually to ensure they met the assumptions of normality and homoscedasticity. When they did not, as was the case for development time, the response variable was power transformed (most suitable transformation determined using the *power.Transform* function in the *car* package of R v3.5.2), and the P-values for this model were obtained using a type 3 Wald Chi-sq. test (function *Anova* in *car* package in R).

We analysed age-independent flying ability of virgin males and females as a binary response (no flight vs. flight). To test binary flight ability, we used a z-test of equality of proportions to compare male and female frequency of flight (function *prop.test* in R package *stats’*). We further tested binary flight ability in virgin males (due to very low virgin female flight responses), using a logistic regression with a binomial distribution (function *glmer* in package *lme4*; Bates et al. 2015), with density as a fixed effect, and container as a random effect.

#### Age-dependent effects

To model age-dependent responses for fecundity, and ability and propensity to fly, we fitted general additive mixed models (function *gam* in package *mgcv*; Wood 2016), with individual identity and container as random effects (using the term bs=“re”). We included adult lifespan as a fixed effect in all models to account for selective disappearance (Pol & Verhulst 2006). For each separate response variable, we started with a global model, containing separate curves for each treatment group. In a first step, we determined the best-fitting smooth classes (thin plate splines, cubic regression splines, shrinkage version of cubic regression splines; tested with standard smooth classes, or tension product smooth classes), using the Akaike Information Criterion (AIC) for model comparisons. The best-fitting smooth classes for the global model was then used in further models.

Flying ability was recorded as an ordinal response, with the first three levels signifying increasing ability to fly (level 1: no wing movement, level 2: wing movement but no lift, level 3: wing movement and lift but no flight), and flight as the fourth level. We therefore used the ordered categorical model family (*ocat*) available for *gam* in R. For this, threshold values of an underlying latent variable (following a logistic distribution) at which the response variable changes, are estimated. To evaluate the effects of variables of interest on the location and shape of age-dependent trajectories, curvature can be modelled as group-specific (e.g. sex-specific) or general (across groups), adding group-specific or general smooth terms to models. If the shape of the estimated trajectory is not different between groups, the trajectory could still differ in location along the y-axis (overall lower or higher values of the response variable). This effect can be modelled by adding a group-specific fixed effect (intercept) to the model (following Zajitschek et al, 2012). We tested the global model against models with either sex-specific curvature or sex-specific curvature and intercept (independent of developmental density), with either density-dependent curvature or density-dependent curvature and intercept (independent of sex), with sex-specific intercept (same curvature, independent of density), and with density-dependent intercept (same curvature, independent of sex). We only used data collected on virgin individuals for this analysis.

To test the effect of mating on flight performance in females, we used a similar modelling strategy, except that we only used female flight measurements recorded between their day of eclosion and 8 days after emergence, because no mated females survived beyond that age. Adult lifespan was not included in the final analyses for all best-fitting models. The AIC of the best fitting model was always at least 2 units smaller than the next best-fitting model (except for effects on earliest age of maximum flight-propensity).

To test whether there were any sex-specific effects of developmental density on age-dependent survival in adult virgin beetles, we modelled the survival data using the non-parametric log-rank test with the *survdiff* function (rho=0) in the *survival* package in R v3.5.2. We tested the effects of density for each sex separately. Kaplan-Meier curves were used to visualise the data. We chose not to use a cox-proportional hazards model because our mortality curves violated the assumption that require the proportional difference in survival between treatments to remain constant over time when checked visually (Hess, 1995).

## Results (Full model outputs in Supplementary Information)

### Age-independent traits

Developmental density had a significant effect on emergence success of beetles with a smaller proportion of beetles emerging from high developmental density than low developmental density containers (P=0.020, z= 2.321, Table S1). Developmental density did not affect sex ratio of eclosed beetles (P=0.626, z= −0.487, Table S2). Developmental time was not affected by an interaction between sex and developmental density (P=0.246, Chisq.= 1.349), or by developmental density (P=0.264, Chisq.= 1.249), but was affected by sex (P<0.001, Chisq.= 43.240), with males developing faster than females (Table S3). Developmental density and sex interacted significantly to affect adult lifespan (P=0.012, DF=1085.474, t= −2.505) indicating that high developmental density reduced female adult lifespan more than it did male adult lifespan. Males developed faster than females (P<0.001, DF=1086.82, t= - 17.734) but density itself had no effect on adult lifespan (P=0.431, DF=22.342, t= −0.802) (Table S4, Figure 1). When looking at the effects of mating on female adult lifespan, we found that mated females lived for a shorter duration than virgin females (P<0.001, DF=577.877, t= 24.830). But female adult lifespan was not affected by an interaction between mating status and developmental density (P=0.250, DF=572.735, t=1.152) or by independent effects of density (P=0.284, DF=53.486, t= 1.081) (Table S5). Female fecundity was affected by developmental density (P<0.001, Chisq.= 11.255), with females from high density treatments laying significantly fewer eggs than females from low developmental density (mean eggs laid ± SE: High density= 75.800 ± 1.700, Low density = 87.900 ± 1.800) (Table S6).

**Figure 1.**
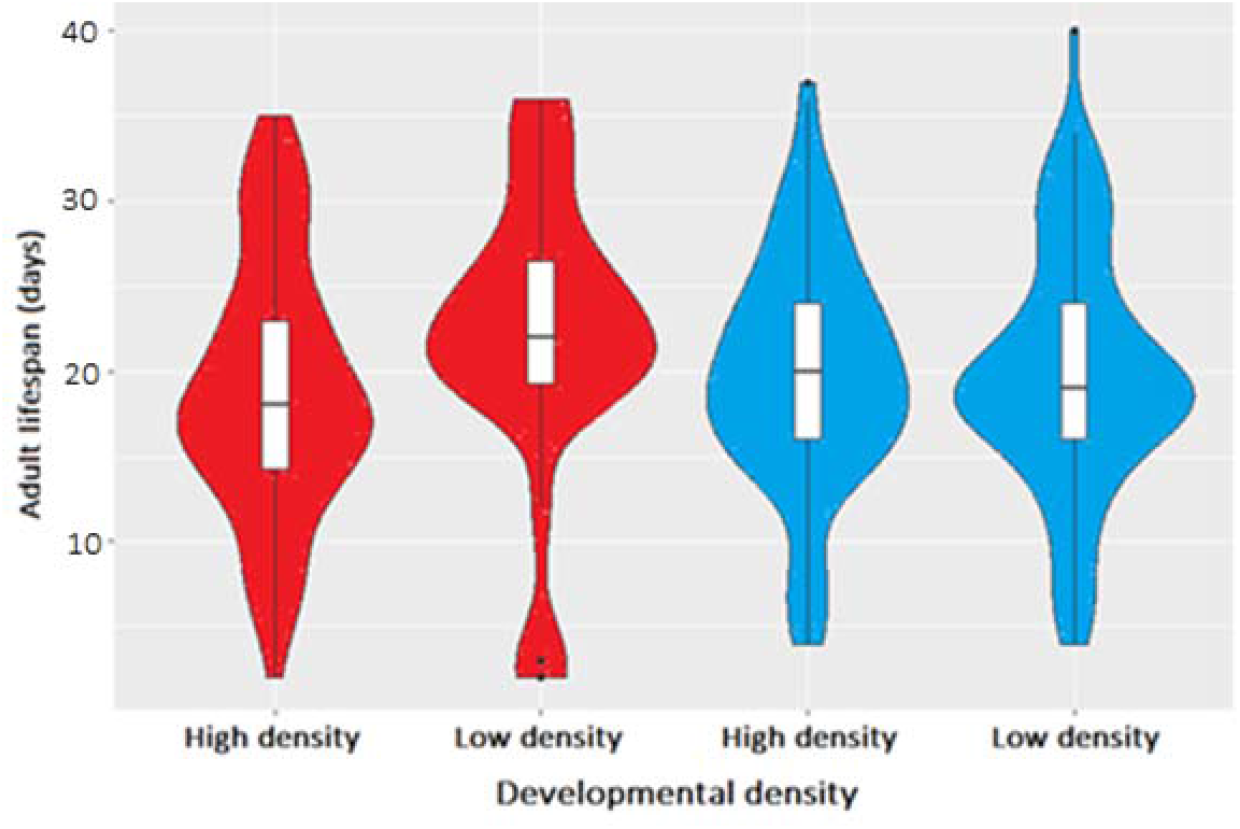
Shows the effect of sex (Blue: male, Red: female) and developmental density (high and low) on adult lifespan. Smoothed probability density of adult lifespan data is represented using violin plots with median and interquartile ranges.

### Age-dependent traits

#### Eggs laid per day

Density had a significant effect on age-dependent fecundity trajectories. A model with density-specific curvature provided the best fit compared to models with either one overall curve (same shape and location), or a model that allowed for a different location along the response variable axis (overall lower and higher fecundity), but with identical overall curvature (Table S7). Low density females laid more eggs in early adult life, compared to high density females, with very similar fecundity being observed in mid- and late-adult life (Figure 2).

**Figure 2.**
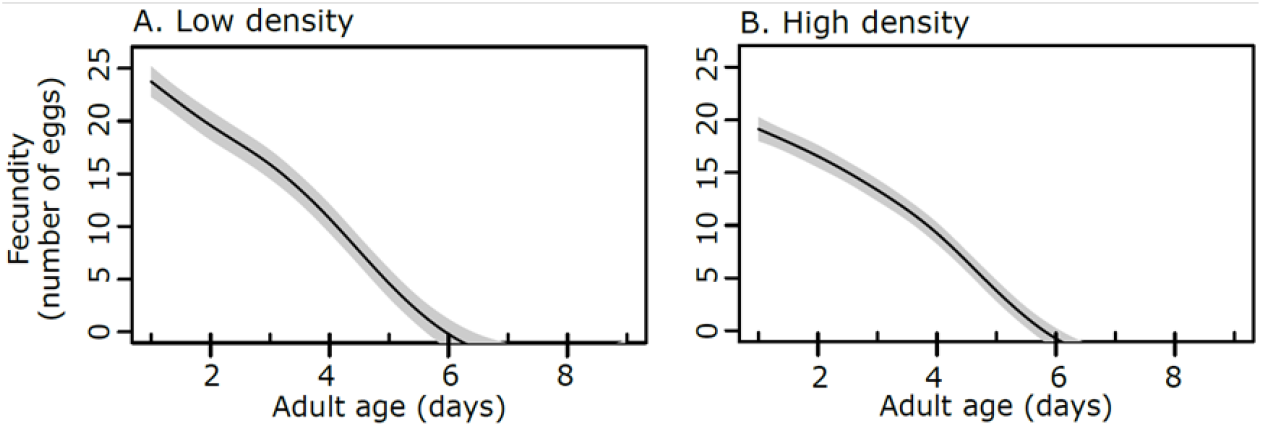
Effects of age (in days) and density on female fecundity for beetles from A.) Low density and B.) High density. Grey shaded bands represent 95% confidence intervals.

We compared virgin male and virgin female age-independent flying ability by assigning them a binary score (flight vs. no flight). Virgin males were more likely to fly than females (binary flying ability, no flight vs. flight, z-test: χ^2^ = 14.030, DF = 1, P< 0.001). Out of 377 measured virgin male (n=259, with 2157 individual measurements) and female (n=118, with 1086 individual measurements) beetles, only 3 females and 43 males flew at least once when physically stimulated. Due to the very low numbers of virgin females exhibiting flight, we only tested whether developmental density had an effect on binary flying ability in virgin males. Density had no effect on binary flying ability in virgin males (GLMM: coef = 0.064, SE = 0.234, *z* = 0.273, P= 0.785).

#### Flying ability

##### Virgin males versus virgin females

The best model for age-dependent flying ability (measured as a 4-level response variable) included sex-specific and density-specific trajectories (Table S8, Figure 3). Males showed overall higher flying ability than females, which was also evidenced by the significant effect of sex when added as a fixed effect in the best-fit model (sex as fixed effect: coef = 1.115, SE = 0.328, *z* = 3.405, P < 0.001). Density had no such effect on age-dependent flying ability (coef = −0.230, SE = 0.328, *z* = −0.702, P= 0.483).

**Figure 3.**
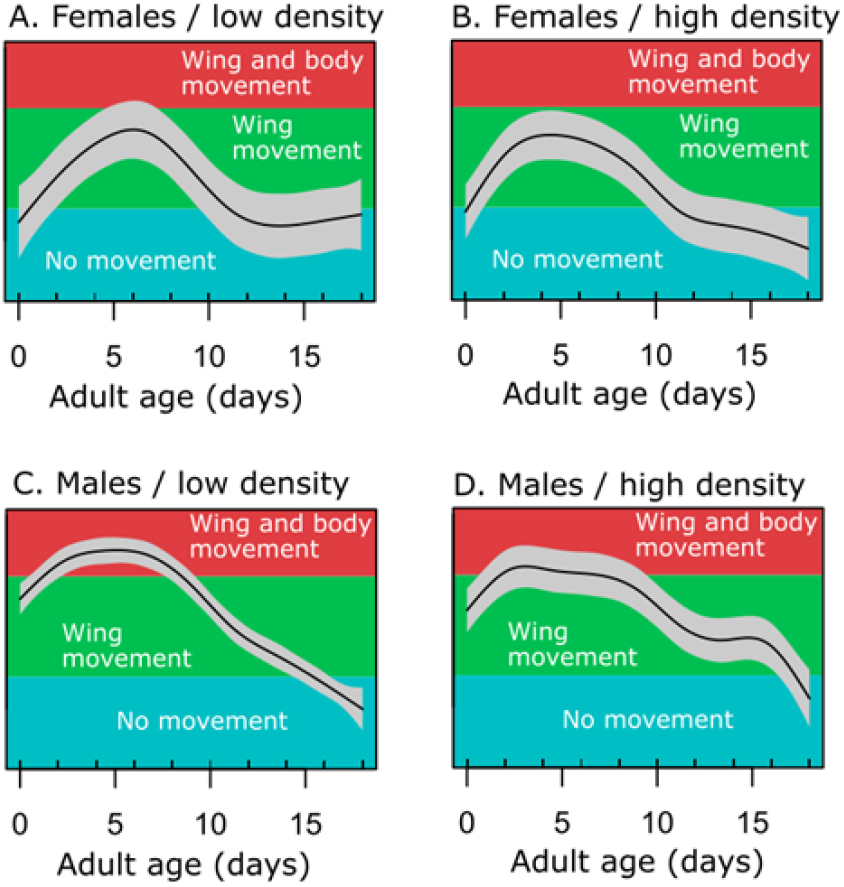
Effects of age, sex, and density on flying ability in virgin female (A, B) and male (C, D) beetles. Flying ability is an ordinal response and each level is represented as a different colour. Flight, the highest level of the *flying ability* variable, is not shown, as the low numbers of recorded instances of flight did not result in a prediction of flight. Grey shaded bands represent 95% confidence intervals.

##### Virgin females versus mated females

Mated females had far fewer measurements of flight taken during their shorter lives (mean ± SE of 3.530 ± 0.070 days, maximum of 5 measurements before death), compared to virgin females (mean ± SE of 9.200 ± 0.180 days, maximum of 10 measurements before death). For this reason, we only used the first 5 flight measurements of virgin and mated females to test for treatment effects on flying ability. The model with two separate terms for virgin and mated females but not for density, was the best fitting model (Table S9). Mated females showed a clear ageing pattern, with a peak in flying ability seen at two days of age followed by a decline over the next three days, with no such pattern in virgin females (Figure 4).

**Figure 4.**
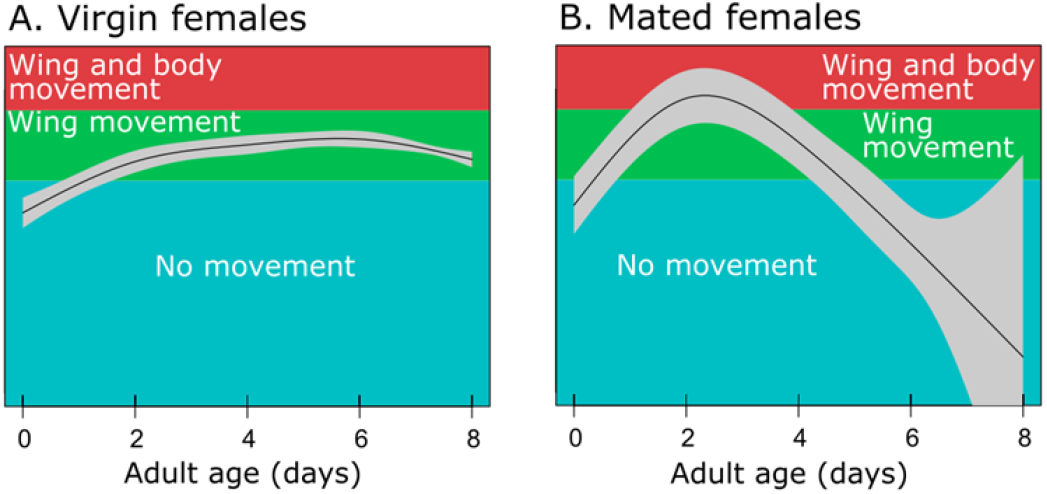
Effect of mating on flying ability for the first 8 days after emergence for A.) virgin and B.) mated females. Flying ability is an ordinal response and each level is represented as a different colour. Flight, the highest level of the *flying ability* variable, is not shown, as the low numbers of recorded instances of flight did not result in a prediction of flight. Grey shaded bands represent 95% confidence intervals.

#### Propensity to fly in virgin beetles

The best model for age-dependent propensity to fly (Table S10) and earliest age of peak in propensity to fly (Table S11), included sex-specific trajectories. Virgin females showed a later and lower peak than virgin males (Figure 5). The first age at which maximum propensity to fly was reached was on average 2.300 ± 0.340 SE days earlier in males compared to females. Density had no effect.

**Figure 5.**
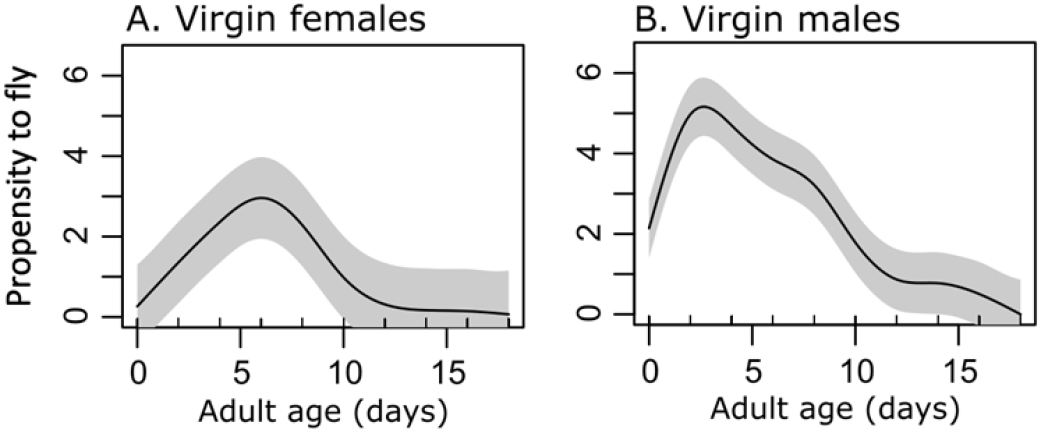
Effect of sex on the propensity to fly for A.) Virgin females and B.) Virgin males with increasing age. Propensity was measured as the number of times a beetle performed it flying ability action. Grey shaded bands represent 95% confidence intervals.

#### Age-dependent survival (Figure 6)

Within virgin males, there was a significant effect of density on age-dependent survival (Chisq.= 8.1, DF= 1, P= 0.004, Table S12a), with males developing in low density showing a steeper decline in age-dependent survival than males from high density. This effect of density on age-dependent survival was not observed in females (Chisq.= 0.5, DF= 1, P=0.472, Table S12b).

**Figure 6.**
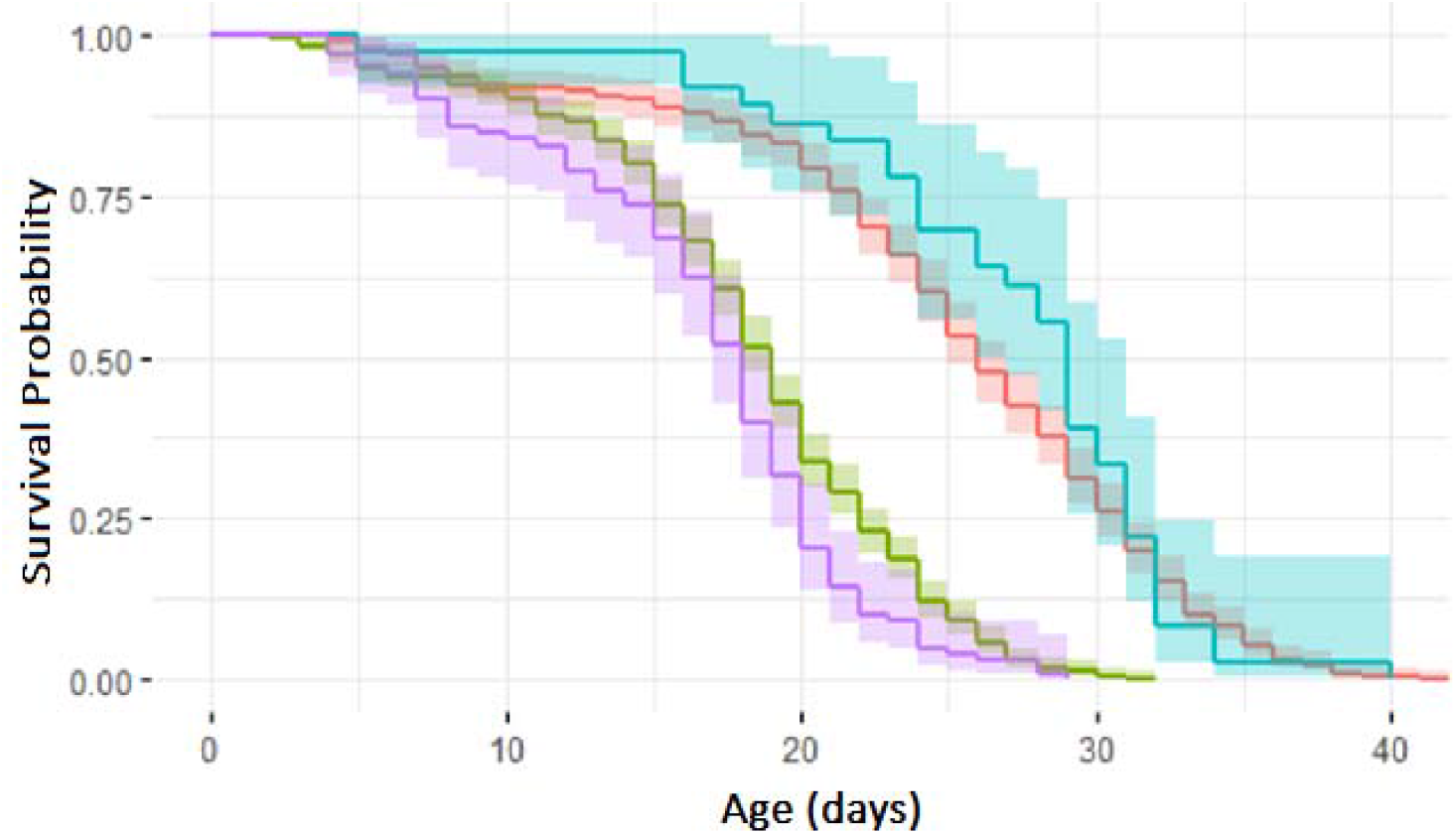
The probability of survival of virgin males and females from high and low developmental densities, with increasing age (time), using Kaplan-Meier curves. Shaded error bars represent 95% confidence intervals. (Purple: males from low density; Green: males from high density; Red: females from high density; Blue: females from low density).

## Discussion

The effects of environmental stress experienced during development can have long lasting consequences for fitness and ecologically relevant traits (Monaghan et al, 2011). Developmental environments can not only affect individuals at a given point in time, but throughout their lives, hence influencing how these traits deteriorate and senesce (Cooper & Kruuk, 2018). To test this, we experimentally manipulated the developmental densities of male and female *C. maculatus*, by subjecting them to either developmental crowding or not. We then measured their resultant developmental (developmental time, egg to adult viability) and adult traits (eggs laid, lifespan, flying ability, and propensity to fly) throughout their adult lives.

Contrary to our expectation, development time was not affected by density or by a density by sex interaction. It has been shown that early life stress, including developmental crowding can decrease development time (Ireland & Turner, 2006; Ludewig et al, 2017). The decrease in development time with higher densities is usually predicted to occur because it allows organisms to emerge sooner to acquire resources and mates earlier than their competitors (Bauerfeind & Fischer, 2005). However, *C. maculatus* is facultatively phytophagous and does not need to feed after emergence in order to survive and reproduce (Credland et al, 1986) which may explain why we did not see high developmental density accelerate development. We did however find that males develop quicker than females, a pattern which has been found previously by Harano & Miyatake (2011), and could be driven by the need for males to compete for access to females. The fact that this pattern is not affected by developmental density or by its interaction with sex, but is affected by sex suggests that it could be sexual selection rather than resource competition which controls seed beetle developmental rate.

Our results showed that sex and developmental density interacted to affect adult lifespan, with female adult lifespan being more negatively affected by high developmental density than male lifespan. One reason for this could be that females of *C. maculatus* are larger than males, hence would require more space and resources inside the bean when developing than males. Consequently, this would mean that the effect of developmental competition would be more severe for females than males as has also been seen in a butterfly (Fantinou et al, 2008). In support of this, previous research on *C. maculatus* by Iglesias-Carrasco et al (2020) has shown that males were generally less affected by larval competition than females. However, our results contrast with studies on other species which have shown developmental crowding (Descamps et al, 2008) and early life stress (Lee et al, 2013) to reduce adult lifespan in both males and females, while Wilkin & Sheldon (2009) show no effect of early-life stress on female lifespan but a significant effect on male lifespan. These differences could be attributed to the different physiologies of the species being studied. We did observe a difference in emergence rates of beetles between high and low developmental density treatments. But because there was no difference in sex ratios of emerged beetles between treatments, it is unlikely that the lack of effect of density on male adult lifespan is due to selective disappearance of males at the developmental stage.

As predicted, we showed that individuals which experienced developmental crowding bore a fitness cost as adults. Specifically, females from high density treatments had a lower total lifetime fecundity than females from low density treatments. This detrimental effect of high density on fecundity has been shown previously in animals such as leaf beetles (Muller et al, 2016), *Drosophila* (Prout & McChesney, 1985), salamanders (Scott, 1994), and in *C. maculatus* (Credland et al, 1986, Vamosi, 2005). There is also strong evidence which suggests that in general, stressful early-life environments should affect fecundity negatively (Hayward et al, 2013; van de Pol et al, 2006). This effect is likely to demonstrate a “silver-spoon effect” which states that individuals which experience optimum early life conditions have an overall higher fitness and performance in their adult lives (Monaghan, 2008, Cooper & Kruuk, 2018) than those who experience stressful environments.

In addition, we found a clear effect of density on age-dependent declines in female fecundity. Females from low density environments showed higher early adult life reproduction than females from high density. This indicates that females from competition-free developmental conditions invest more in early adult life reproduction than those from poor developmental environments possibly due to having a better body condition and a greater number of eggs. Higher investment in early-life reproduction has been shown to cause a faster rate of reproductive senescence (Bowhuis et al, 2010; Hammers et al, 2013) because of hypothesized trade-offs between early-life reproduction with late-life reproduction and survival, although surprisingly we did not observe this. A possible reason for this could be that females developing in low densities are in much better condition than females from high densities. This could allow females from low developmental densities to reproduce more and live longer (Reznick et al, 2000), without experiencing accelerated deterioration in reproduction with age that is caused by a lifespan-reproduction trade-off.

In males, developmental density affected age-dependent survival, with males from low density treatments showing steeper decline in survival with increasing age than males from high density treatments. One reason for this could be that low density males invest more in reproductive traits such as sperm production, leading to trade-offs between reproduction and age-dependant survival. This is something future studies could investigate. No effect of developmental density was observed on age-dependant survival in females. This reinforces our previously stated finding that females from low developmental density may not bear a cost of higher early life investment in reproduction, on survival, due to being of superior condition than females from high developmental densities. These sex-specific effects on age-dependent survival have also been shown by Zajitschek et al (2009) in crickets, and point towards different life history strategies of males and females.

Developmental density did not influence flight performance of either sex. Instead significant effects of sex show that males generally flew better and more often than females, and that male flying ability and propensity peaked higher and at an earlier age than females. One reason for males being more willing to fly could be due to sex biased dispersal (Pusey, 1987). Sex-biased dispersal arises due to different costs (energetic, predation) and benefits (novel environment, mating opportunity, inbreeding avoidance, reduced competition) of dispersal to each sex (Baines et al, 2017, Li & Kokko, 2019). Although most evidence for sex-biased dispersal comes from mammals and birds (Li & Kokko, 2019), laboratory studies such as Downey et al (2015) and Mishra et al (2020), which measured dispersal as crawling distance, have shown male-biased dispersal in *Callosobruchus* and *Drosophila* respectively. A recent study also showed male biased dispersal in butterflies during the early breeding season, but female biased dispersal during the late season (Plazio et al 2020). The authors suggested that these differences were due to different reproductive strategies of males and females, which could also be the case in *C. maculatus*.

Our study showed a difference in age-dependent trajectories in flight ability between mated versus unmated females, with mated females showing a higher ability to fly as well as a steeper decline in flying ability with age. Additionally, mated females also had a reduced lifespan compared to virgin females. The difference in flight between virgins and mated females may arise because females try to avoid male harassment and mating costs after having mated once, due to additional mating with the same male providing no benefit to females (Okada et al, 2019). It could also be because females want to disperse after having mated, in search of beans to lay their eggs on. These results are consistent with costs of mating and male-harassment associated with other female traits such as adult lifespan and fitness (den Hollander & Gwynn, 2009; Gay et al, 2009; Ronn et al, 2006).

For virgin individuals of both sexes, flight was affected by age, with flying ability and propensity to fly increasing in early-adult life and then deteriorating in late-adult life. This pattern of initial increase followed by deterioration with age follows a classic senescence curve (Jones et al, 2014), and has been reported by Ahman & Karlsson (2009) in the flight of butterflies. Such age-related changes could be attributed to changes in flight muscles with age, as shown in *Drosophila* (Miller et al, 2008). Increase in the accumulation of free radicals with age due to the metabolic costs associated with flying could also explain why flight traits deteriorate with age (Lane et al, 2014). Though our definition of senescence has been limited mostly to classic fitness components, performance traits like flight, which could affect processes such as mate finding, dispersal, and hence fitness, can also change and deteriorate with age. Thus, greater consideration of the age-dependence of flight (or indeed other forms of dispersal), is important to help us understand ecological patterns and processes such as age-structure in, and age-dependent gene flow between populations.

Density-dependent development of dispersal morphs has been predicted to arise when individuals need to disperse from crowded conditions to avoid competition and has been shown in grasshoppers, water striders, and aphids (Applebaum & Heifetz, 1999; Denno & Roderick, 1992; Han, 2020; Harrison 1980). In these species, individuals that were raised in high densities develop a dispersal morph while individuals raised in low densities develop a sedentary morph. Our experiment, which unlike previous studies on *Callosobruchus* (Appleby & Credland, 2001; Messina & Renwick, 1985; Utida, 1972), measures flying ability throughout an individual’s life and on both sexes using a repeatable method, did not find that crowded developmental conditions induced higher flight, hence dispersal ability. But we did find significant effects of density on other life history traits in females such as lifespan, reproductive senescence, and total fecundity. Although these results do not support presence of density-dependent dispersal morphs, they do demonstrate density-dependent differences in life history traits.

## Conclusion

Our study substantiates previous findings of how the developmental environment affects life history and performance traits. It is also one of the first to test the effects of developmental environment on age-dependent flight in both males and females. We show that although developmental crowding has no effect on male lifespan or on age-dependent flight performance of both sexes, it has a significant effect on female fitness traits of fecundity, reproductive senescence, and lifespan, as well as on male age-dependent survival. We also show strong sex-specific differences in male and female flight, which may be influenced by male and female dispersal decisions and mating strategies. Overall, our results can be summarised as two findings. First, there is a “silver-spoon” effect of developmental environments on female fitness components and developing in low density conditions is beneficial only to females, with no such benefits being observed in males. Second, the differences in male and female age-dependent flight reflect differences in male and female behaviour, physiology, and life history strategy. We suggest that males have evolved to prioritise dispersal and outbreeding, which they do by flying more and earlier than females, whereas females have evolved to prioritise mating, and once mated, avoid additional costs of mating by flying away to search for beans to lay eggs on. Our study has implications for understanding not only the effects of developmental environments on age-dependent changes in traits, but also the population-level and ecological changes brought about by sex-specific responses to the developmental environment at different ages.

