## Supplementary material for "Sex-specific effects of developmental density on life history and age-dependent flight traits"

**Appendix S1**

Repeatability and comparison of methods used to measure flight

To measure repeatability of our flight performance measures, we conducted a pilot test on a stock population of beetles before the start of our experiment. 50 females and 104 males were chosen randomly and measured independently by two observers, each using two different methods, on the same day in a randomised order. The observers were unaware of each other’s scores.

The first method has been described in the main text of our paper, and uses forceps to tap beetles on their pygidium to elicit a flight response, and then given them an ordinal score. Using this method, 70% of data points showed the same values for the two observations, when measured using an ordinal score from “0” to “3” recorded by tapping on individuals. Just by chance, the probability of getting the same score would be 25% for each observation. Out of all the beetles which were recorded to fly by either observer, 81.8% of individuals were recorded to fly by both (‘3’ score). Similarly, out of all the beetles recorded to show wing and body movement by either observer, 60.7% of individuals were recorded to show wing and body movement by both observers (‘2’ score). Likewise, 44.8% of individuals were recorded to show only wing movement by both observers, while the rest 55.2% by only one observer (‘1’ score). And 44.1% of individuals were recorded to show no movement (‘0’ score) by both observers out of all the beetles which were recorded with a “0” score. *Supplementary videos which show examples of the different ordinal scores of beetle flight ability have been uploaded under an anonymous file on Open Science Framework and can be accessed using this link:* [*https://osf.io/2dp8r/?view_only=0e428ce25c284db195972c4b01e53057*](https://osf.io/2dp8r/?view_only=0e428ce25c284db195972c4b01e53057) *.*

To compare this method to another method previously used by Utida (1972) and Appleby & Credland (2001) we conducted another experiment on the same subset of beetles using Appleby & Credland (2001)’s described method. In this case, the beetles were lifted to a height of 50-100 cm and dropped. They were scored as either “1” if they flew each time, or “0” if they did not fly. This process was repeated by another observer on the same individual, and the order of observers taking turns was randomised.

This alternative method for measuring flight showed a repeatability of 88% between observers. Just by chance, the probability of getting the same score would be 50% for each observation. Only 7% of beetles were recorded by both observers as fliers (“1”) while 82% of beetles were recorded by both observers as non-fliers (“0”).

Because the method of tapping the individual gave a more nuanced score, elicited a flight response more often in beetles hence reducing false-negatives for ability to fly, and had a greater ratio of repeatability of flight to that expected by chance than the method which dropped beetles, we chose to use the former.

**Appendix S2**

**Table S1.** Effect of **developmental density on emergence success** of beetles. Results were obtained using a logistical regression model using the function *glm* in package *stats* in R and modelling emergence as a binary variable (“logit” link function used: 1 for emerged or 0 for did not emerge). Significant values are highlighted in bold. Container was not used as a random effect because container was our level of replication.

| **Term** | **Estimate** | **SE** | **z** | **P** |
| --- | --- | --- | --- | --- |
| Intercept | 1.980 | 0.089 | 22.273 | <0.001 |
| Density (Low) | 0.697 | 0.300 | 2.321 | **0.020** |

**Table S2.** Effect of **developmental density on sex ratio** of emerging beetles. Results were obtained using a logistical regression model using the function *glm* in package *stats* in R and modelling emergence as a binary variable (“logit” link function used). Container was not used as a random effect because container was our level of replication.

| **Term** | **Estimate** | | **SE** | | **z** | | **P** |
| --- | --- | --- | --- | --- | --- | --- | --- |
| Intercept | 0.061 | 0.062 | | -0.993 | | 0.321 | |
| Density (Low) | 0.078 | 0.159 | | -0.487 | | 0.626 | |

**Table S3.** Effect of **density and sex on development time**. Development time was power transformed by x^-7 using the function *powerTransform* in R package *car*, to meet the assumption of normality. Because of a negative power transform, estimates should be interpreted as inverted in their sign. P-values obtained using type 3 Wald chi sq. test using the function *Anova* in R. “Full model” provide parameter estimates and significance values for interpreting the interaction only (in grey), while main-effects model do not fit interactions, and provide parameter estimates only to interpret main-effects (in grey). Model was run as an LMM conducted using package *lme4* in R. Significant values are highlighted in bold.

| **Model** | **Term** | **Estimate  x 10^-11** | **SE x 10^-11** | **χ2** | **P** |
| --- | --- | --- | --- | --- | --- |
| Full | Fixed effects |  |  |  |  |
|  | (Intercept) | 17.380 | 0.719 | 584.095 | <0.001 |
|  | Density (Low) | -1.669 | 1.130 | 2.181 | 0.140 |
|  | Sex (Male) | 1.758 | 0.315 | 31.398 | <0.001 |
|  | Density*Sex | 0.935 | 0.805 | 1.349 | 0.246 |
|  | **Random effects** | **Variance x 10^-11** | **SD x 10^-11** |  |  |
|  | Container | 2.33E-10 | 1.53 |  |  |
|  | **Term** | **Estimate  x 10^-11** | **SE x 10^-11** | **χ2** | **P** |
| Main effect | Fixed effects |  |  |  |  |
|  | (Intercept) | 17.310 | 0.718 | 580.294 | <0.001 |
|  | Density (Low) | -1.172 | 1.049 | 1.249 | 0.264 |
|  | Sex (Male) | 1.900 | 0.289 | 43.240 | **<0.001** |
|  | **Random effects** | **Variance x 10^-11** | **SD x 10^-11** |  |  |
|  | Container | 2.35E-10 | 1.53 |  |  |

**Table S4.** Effect of **density and sex on lifespan** of **virgin beetles**. “Full model” provide parameter estimates and significance values for interpreting the interaction only (in grey), while main-effects model do not fit interactions, and provide parameter estimates only to interpret main-effects (in grey). Model was run as an LMM using package *lme4* in R. Significant values are highlighted in bold.

| **Model** | **Terms** | **Estimate** | **SE** | **DF** | **t** | **P** |
| --- | --- | --- | --- | --- | --- | --- |
| Full | Fixed effects |  |  |  |  |  |
|  | (Intercept) | 25.178 | 0.427 | 10.865 | 58.992 | <0.001 |
|  | Sex (Male) | -6.962 | 0.433 | 1085.254 | -16.072 | <0.001 |
|  | Density (Low) | 1.858 | 1.223 | 154.559 | 1.519 | 1.307 |
|  | Sex (Male)  * density (Low) | -3.433 | 1.37 | 1085.474 | -2.505 | **0.012** |
|  | **Random effects** | **Variance** | **SD** |  |  |  |
|  | Container (intercept) | 0.389 | 0.624 |  |  |  |
|  | **Terms** | **Estimate** | **SE** | **DF** | **t** | **P** |
| Main  effects | Fixed effects |  |  |  |  |  |
|  | (Intercept) | 25.369 | 0.426 | 9.792 | 59.567 | <0.001 |
|  | Sex (Male) | -7.306 | 0.412 | 1086.82 | -17.734 | **<0.001** |
|  | Density (Low) | -0.594 | 0.741 | 22.342 | -0.802 | 0.431 |
|  | **Random effects** | **Variance** | **SD** |  |  |  |
|  | Container (intercept) | 0.412 | 0.642 |  |  |  |

**Table S5.** Effect of **mating status of females on female adult lifespan**. “Full model” provide parameter estimates and significance values for interpreting the interaction only (in grey), while main-effects model do not fit interactions, and provide parameter estimates only to interpret main-effects (in grey). Model was run as an LMM using package *lme4* in R. Significant values are highlighted in bold.

| **Model** | **Terms** | **Estimate** | **SE** | **DF** | **t** | **P** |
| --- | --- | --- | --- | --- | --- | --- |
| Full | Fixed effects |  |  |  |  |  |
|  | Intercept | 6.428 | 0.847 | 150.194 | 7.586 | <0.001 |
|  | Status (virgin) | 18.712 | 0.901 | 576.689 | 20.763 | <0.001 |
|  | Treatment (Low) | -0.119 | 1.307 | 182.102 | -0.091 | 0.927 |
|  | Status (virgin) *  Treatment (Low) | 2.033 | 1.764 | 572.735 | 1.152 | 0.250 |
|  | **Random effects** | **Variance** | **SD** |  |  |  |
|  | Container | 0.088 | 0.298 |  |  |  |
|  | **Terms** | **Estimate** | **SE** | **DF** | **t** | **P** |
| Single  effects | Fixed effects |  |  |  |  |  |
|  | Intercept | 5.969 | 0.751 | 93.328 | 7.949 | <0.001 |
|  | Status (virgin) | 19.243 | 0.775 | 577.877 | 24.83 | **<0.001** |
|  | Treatment (Low) | 0.975 | 0.902 | 53.485 | 1.081 | 0.284 |
|  | **Random effects** | **Variance** | **SD** |  |  |  |
|  | Container | 0.104 | 0.323 |  |  |  |

**Table S6.** Effect of **developmental density on lifetime female fecundity**. Results are obtained by running an LMM using package *lme4* in R. Significant values are highlighted in bold.

| **Terms** | **Estimate** | **SE** | **χ^2^** | **P** |
| --- | --- | --- | --- | --- |
| Fixed effects |  |  |  |  |
| (Intercept) | 75.753 | 2.422 | 978.08 | <0.001 |
| Density (Low) | 12.07 | 3.598 | 11.255 | **<0.001** |
| **Random effects** | **Variance** | **SD** |  |  |
| Container | 16.18 | 4.022 |  |  |

**Table S7**. Effects on **age-dependent** **fecundity**. Comparison of three models with either a separate smoothing term (curve) for high and low density (Density smooths), one overall smoothing term (Overall smooth), or identical smooth for low and high developmental densities, allowing a density specific intercept (overall height of the curve: Overall smooth and density intercept). Models constructed using GAM smooth classes: tensor product smoothers te(), penalized cubic regression splines (cr). Additional effects in all models: lifespan (fixed effect), container (random effect), individual ID (random effect). df: degrees of freedom. dAIC: difference in AIC to the best-fitting model.

| Model | Smooth terms | Fixed effect | df | dAIC |
| --- | --- | --- | --- | --- |
| Density smooths | 2 (low, high density) |  | 61.14 | 0.0 |
| Overall smooth | 1 (overall) |  | 65.52 | 27.5 |
| Overall smooth and  density intercept | 1 (overall) | density | 54.80 | 28.0 |

**Table S8**. Effects on **age-dependent flying ability**. The most complex tested model (Mating x density smooths) fits a separate curve to each of the four treatment groups (sex x density) and provides the best fit. Models constructed using GAM smooth classes: tensor product smoothers te(), penalized cubic regression splines (cr). Additional effects in all models: lifespan (fixed effect), container (random effect), individual ID (random effect). df: degrees of freedom. dAIC: difference in AIC to the best-fitting model.

| Model | Smooth terms | Fixed effect | df | dAIC |
| --- | --- | --- | --- | --- |
| Mating x density smooths | 4 (virgin/low, virgin/high, mated/low, mated/high) |  | 324.79 | 0.0 |
| Sex smooths | 2 (male, female) |  | 309.17 | 8.7 |
| Overall smooth and  sex intercept | 1 (overall) | sex | 303.31 | 30.1 |
| Density smooths | 2 (low, high density) |  | 318.46 | 34.8 |
| Overall smooth | 1 (overall) |  | 312.22 | 42.7 |
| Overall smooth and  density intercept | 1 (overall) | density | 312.86 | 43.6 |

**Table S9**. Effects on **age dependent flying ability** in **virgin and mated female** beetles. The model with only mating status (Mating smooths) provided the best fit when compared to other models. Models constructed using GAM smooth classes: tensor product smoothers te(), penalized cubic regression splines (cr). Additional effects in all models: lifespan (fixed effect), container (random effect), individual ID (random effect). df: degrees of freedom. dAIC: difference in AIC to the best-fitting model.

| Model | Smooth terms | Fixed effect | df | dAIC |
| --- | --- | --- | --- | --- |
| Mating smooths | 2 (virgin, mated) |  | 149.23 | 0.0 |
| Mating x density smooths | 4 (virgin/low, virgin/high, mated/low, mated/high) |  | 156.68 | 2.3 |
| Overall smooth | 1 (overall) |  | 132.62 | 157.3 |
| Overall smooth and  density intercept | 1 (overall) | density | 132.84 | 157.7 |
| Overall smooth and  mating intercept | 1 (overall) | mating | 133.17 | 158.2 |
| Density smooths | 2 (low, high density) |  | 135.89 | 160.2 |

**Table S10**. Effects on **age-dependent propensity to fly**. The model in which only includes sex as a smooth provides the best fit when compared to models with either an interaction of mating status and density (Mating x density smooths), or overall smooth with a sex intercept and sex or density as a fixed effect, or only density smooths, or only overall smooths. Models constructed using GAM smooth classes: standard smoothers s(), thin plate splines (tp). Additional effects in all models: lifespan (fixed effect), container (random effect), individual ID (random effect). df: degrees of freedom. dAIC: difference in AIC to the best-fitting model.

| Model | Smooth terms | Fixed effect | df | dAIC |
| --- | --- | --- | --- | --- |
| Sex smooths | 2 (male, female) |  | 301.85 | 0.0 |
| Mating x density smooths | 4 (virgin/low, virgin/high, mated/low, mated/high) |  | 312.93 | 5.1 |
| Overall smooth and  sex intercept | 1 (overall) | sex | 292.33 | 118.3 |
| Density smooths | 2 (low, high density) |  | 304.54 | 122.7 |
| Overall smooth | 1 (overall) |  | 297.96 | 123.4 |
| Overall smooth and  sex intercept | 1 (overall) | density | 298.06 | 123.8 |

**Table S11**. Effects on **earliest age of maximum propensity to fly**. The model with only sex as a fixed term provides the best fit compared to models with sex and density as either additive or interactive terms, or a model with just the effect of density. Models constructed using Linear mixed model (*lmer* in R). Additional effects in all models: container (random effect). df: degrees of freedom. dAIC: difference in AIC to the best-fitting model

| Model terms (fixed) | df | dAIC |
| --- | --- | --- |
| Sex | 4 | 0.0 |
| Sex + density | 5 | 1.0 |
| Sex + density + sex x density | 6 | 3.0 |
| Density | 4 | 40.5 |
| (intercept only) | 3 | 43.3 |

**Table S12.** Overall effects of a.) developmental density, b.) Sex, as well as c.) and d.) sex-specific effects of developmental density on **adult age-dependent survival** using log-rank test. Models constructed using *survdiff(rho=0)* function in the package *survival* in R. Significant values highlighted in bold.

a. Effect of density on age-dependent survival in males.

| **Density** | **N** | **Observed** | **Expected** | **(O-E)^2/E** | **(O-E)^2/V** |
| --- | --- | --- | --- | --- | --- |
| High | 530 | 530 | 551.541 | 0.841 | 8.105 |
| Low | 98 | 98 | 76.459 | 6.069 | 8.105 |
| **Term** | **Chisq.** | **DF** | **P** |  |  |
| Density | 8.1 | 1 | **0.004** |  |  |

b. Effect of density on age-dependent survival in females

| **Density** | **N** | **Observed** | **Expected** | **(O-E)^2/E** | **(O-E)^2/V** |
| --- | --- | --- | --- | --- | --- |
| High | 427 | 427 | 422.943 | 0.039 | 0.518 |
| Low | 36 | 36 | 40.057 | 0.411 | 0.518 |
| **Term** | **Chisq.** | **DF** | **P** |  |  |
| Density | 0.5 | 1 | 0.472 |  |  |
